# Mouse Neural Organoids Model the Mature *In Vivo* Synaptic Proteome

**DOI:** 10.64898/2026.09.24.753785

**Authors:** Stefano L. Giandomenico, Marc van Oostrum, Quinn Waselenchuk, Irene Dalla Costa, Julian D. Langer, Erin M. Schuman

## Abstract

Neural organoids represent a promising approach for modelling neural circuits and neurological disease, yet their utility depends on whether they can recapitulate the molecular complexity of mature brain synapses. Here, we optimized a protocol to generate mature forebrain organoids from mouse embryonic stem cells and used time-resolved proteomics to define the emergence of synaptic proteins during maturation. Using fluorescence-activated synaptosome sorting coupled with mass spectrometry, we quantified the synapse-enriched proteome and found a high correlation with matched *in vivo* synaptic proteomes. These findings demonstrate that complex mature synaptic molecular architecture is present within an *in vitro* organoid environment.

## Introduction

Neural organoids have recently emerged as powerful *in vitro* systems for studying key aspects of brain tissue development and function in a controlled environment, while retaining elements of three-dimensional architecture and cellular diversity^1–7^. Organoids are capable of generating mature neurons that fire action potentials and establish network activity, underscoring their potential to model early stages of neural circuit formation^2,3,5,8^.

The ability to generate neural organoids from human pluripotent stem cells (PSCs) offers the prospect of studying neurological disease in a human-relevant context^9–14^, but whether these systems can faithfully and fully model the synapse remains an open question. Because proteins are the primary executors of cellular function, and because alterations in synaptic proteins are a hallmark of many neurological disorders^15–18^, it is essential to establish whether *in vitro* 3D neural systems can recapitulate the complex molecular composition of synapses^19–25^ or whether intrinsic limitations of differentiation and culture conditions constrain the assembly of this architecture^26–28^.

Determining the fidelity of organoid-derived synapses requires a comprehensive molecular characterisation and comparison with mature synapses *in vivo*. Recent work addressing this question characterised the composition of growth cone particles and synapses in human organoids, revealing notable similarities to their *in vivo* counterparts^29^. However, the synapses analysed in this study remained at relatively immature developmental stages, leaving unresolved whether their incomplete maturation reflects the intrinsically protracted developmental tempo of human neural systems^26,30–35^ or whether *in vitro* 3D culture imposes fundamental constraints on the acquisition of an *in vivo*-like synaptic molecular composition. Importantly, resolving whether organoid-derived synapses can faithfully reproduce mature synaptic molecular architecture requires both an organoid system that reaches advanced stages of synaptic maturation and well-defined *in vivo* reference datasets of mature synapses.

To answer this question, we developed a protocol to generate mature forebrain organoids from mouse embryonic stem cells^36–38^ and used fluorescence-activated synaptosomes sorting (FASS) and mass spectrometry to compare their synapse-enriched proteomes to matched mature *in vivo* synapses^25,39,40^. We found that synaptic protein enrichment profiles showed a strong correlation between organoid-derived and *in vivo* synapses, demonstrating that *in vitro* organoid systems are capable of establishing the complex molecular organisation characteristic of mature brain synapses.

## Results

To determine whether *in vitro* 3D neural systems can recapitulate the molecular architecture of mature brain synapses, we first established a neural organoid differentiation paradigm capable of supporting advanced stages of synaptic maturation^2,37,41^. Starting from feeder-free mouse embryonic stem cells (mESCs), we developed an unguided forebrain organoid protocol and optimised the timing of media transitions to promote correct tissue identity and long-term maturation. Using this approach, we successfully generated forebrain organoids over a 35-day period (Fig. 1A). The resulting organoids exhibited characteristic neural ventricle-like structures and expressed key markers of neural progenitors (Pax6) and mature neurons (Map2, HuC/D), as well as markers of forebrain divisions (Foxg1), the telencephalon (Tbr2, Emx1, Dlx2) and diencephalon (Gbx2), as assessed by immunolabelling (Fig. 1B and Fig. S1A). By day 18, we observed Vglut1 and Vgat puncta, as well as the post-synaptic scaffold proteins Psd95 and Gphn, and the pre-synaptic marker Bassoon (Bsn), indicating the formation of excitatory and inhibitory synaptic networks (Fig. 1B). Functional assessment by calcium imaging at day 35 confirmed that organoid neurons fire synchronised action potentials that were abolished by tetrodotoxin (TTX) treatment, demonstrating the presence of active, functional synaptic connections (Fig. 1D).

**Figure 1.**
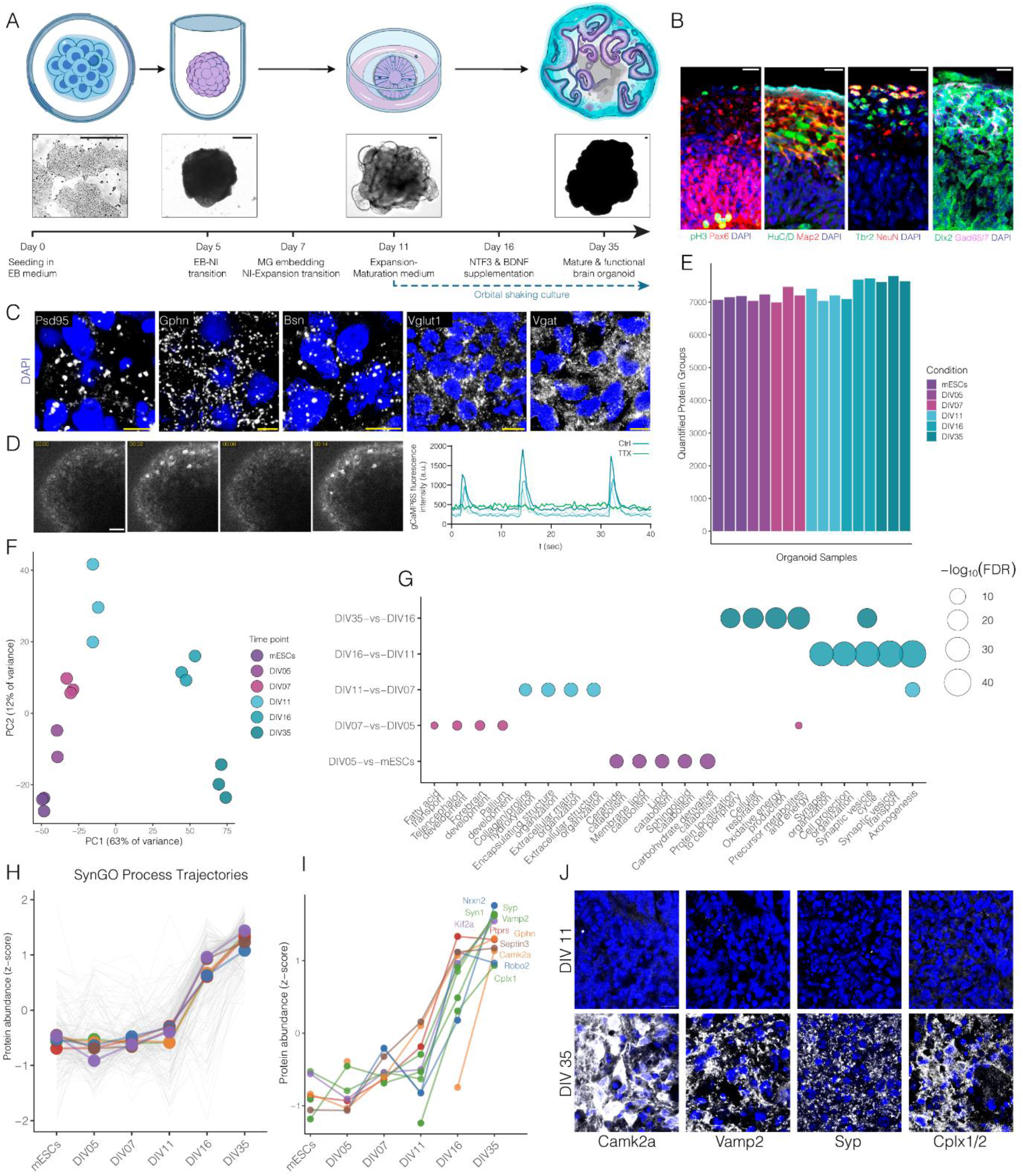
Proteomic profiling captures the developmental trajectory and synaptic maturation of mouse neural organoids. (A) Overview of the mouse brain organoid differentiation workflow. Mouse embryonic stem cells were aggregated to form embryoid bodies (EBs), transitioned through neural induction (NI), expansion medium supplemented with dissolved matrigel, maturation medium, and cultured long-term with NT3 and BDNF supplementation to generate mature brain organoids. Representative bright-field images illustrate organoid size and morphology at key developmental stages (day 0, 5, 11, and 35). Timeline indicates media transitions and major developmental milestones assayed in the proteomics timecourse. Scale bars, 100 μm. (B) Immunofluorescence analysis demonstrating tissue organization, regional identity and neural lineage specification organoids at day 11. Representative confocal images show ventricular zone-like structures and neuronal differentiation marked by pH3 (proliferating progenitors), Pax6 (neural progenitors), HuC/D and Map2 (neurons), Tbr2 (dorsal telencephalon intermediate progenitors), NeuN (mature neurons), and the ventral telencephalon progenitor and interneuron marker Dlx2 together with the GABAergic neuron marker GAD65/67. Nuclei were counterstained with DAPI. Scale bars, 20 μm. (C) Representative immunostaining (day 18 organoids) for synaptic proteins showing expression of the postsynaptic density protein PSD95, the inhibitory postsynaptic scaffold Gephyrin (Gphn), the presynaptic active zone protein Bassoon (Bsn), and excitatory (VGLUT1) and inhibitory (VGAT) vesicular neurotransmitter transporters, demonstrating the establishment of excitatory and inhibitory synaptic compartments. Scale bars, 5 μm. (D) Calcium imaging of day 35 organoids reveals spontaneous network activity. Representative time-lapse images show dynamic calcium transients, and corresponding fluorescence intensity traces demonstrate spontaneous oscillatory activity that is suppressed by tetrodotoxin (TTX), consistent with action potential-dependent neuronal network activity. Each trace corresponds to a recorded neuron. Scale bar, 50 μm. (E) Quantitative proteomic profiling across differentiation. Bar plot shows the number of quantified protein groups detected in each sample from pluripotent stem cells (mESCs) and organoids collected at DIV5, DIV7, DIV11, DIV16, and DIV35, demonstrating high proteome coverage and reproducibility throughout differentiation. (F) Principal component analysis (PCA) of the organoid proteomics time course showing progressive separation of samples according to developmental stage. Replicates cluster tightly within each time point, indicating high reproducibility and temporal proteomic remodeling during organoid maturation. (G) Gene Ontology (GO) enrichment analysis of differentially abundant proteins between consecutive developmental stages. Bubble size represents enrichment significance (−log_10_FDR), and colored bubbles denote the corresponding developmental comparison. Early transitions are enriched for processes associated with neural development, cell proliferation, and metabolism, whereas later stages are characterized by increasing enrichment of synaptic organization, synaptic vesicle cycling, neurotransmission, and axonogenesis. (H) SynGO pathway trajectory analysis illustrating coordinated temporal regulation of synaptic protein abundance throughout organoid maturation. Grey lines represent individual SynGO-associated proteins, whereas colored lines indicate the average trajectory for each developmental stage, demonstrating a marked increase in synaptic protein abundance during late maturation. (I) Representative trajectories of selected presynaptic and postsynaptic proteins, including Nrxn2, Syp, Vamp2, Ptprs, Gphn, Septin3, Camk2a, Robo2, and Cplx1, highlighting coordinated upregulation of proteins involved in synaptic transmission and neuronal connectivity during organoid development. (J) Representative immunofluorescence images validating proteomic findings by demonstrating increased expression of Camk2a, Vamp2, Syp, and Cplx1/2 in mature (DIV35) compared with immature (DIV11) organoids. Nuclei were counterstained with DAPI. Scale bars, as indicated in each panel. Data are representative of independent biological replicates. Scale bars, 10 μm.

Next, to characterise neural organoid development in an unbiased manner and map the trajectories of neuronal maturation and synaptogenesis, we performed proteomic profiling across key developmental stages of the organoid differentiation protocol (Fig. 1A). Proteomic profiling yielded deep coverage across all samples, with over 7,000 protein groups consistently quantified at each developmental stage (Fig. 1E). PCA revealed a clear temporal organisation of the dataset, with samples segregating according to developmental age and replicates clustering tightly together (Fig. 1F). Differential protein expression analyses between successive developmental stages demonstrated a progressive shift from developmental programmes to neuronal maturation, with early enrichment of neurodevelopmental processes followed by strong enrichment of synaptic organisation, synaptic vesicle cycling and axonogenesis at later stages (Fig. 1G). Consistent with a predominantly telencephalic identity, dorsal forebrain markers including Hes1, Tf7l1, Dmrta2, Tbr1, Bbcl11b and Reln were detected throughout differentiation, with deep-layer cortical neuronal markers increasing progressively as development proceeded (Fig. S1B). Ventral telencephalic markers, including Sox6, Meis2, Sp8 and Dlx2, also accumulated over time, indicating the emergence of inhibitory interneuron lineages alongside cortical excitatory neurons (Fig. S1C). In contrast, markers of non-telencephalic fates remained limited, with the thalamic determinant Gbx2 restricted to discrete regions by immunostaining (Fig. S1A), while Otx2 declined following early differentiation and the thalamic marker Hcn4^42^ was detected only at late stages (Fig. S1D), suggesting that diencephalic/thalamic-like populations comprise a minor component of the cultures.

SynGO analysis further demonstrated a progressive expansion of the synaptic proteome during organoid maturation, with proteins significantly increased at DIV16 and 35 showing significant enrichment of both pre- and postsynaptic compartments as well as synaptic signalling, organisation, transport and metabolism (Fig. S1E). Quantification of significantly regulated SynGO proteins revealed that these changes occurred predominantly during the transition from DIV11 to DIV16, with additional increases persisting through DIV35 across multiple functional synaptic categories (Fig. 1H and Fig. S1F). Consistent with this coordinated synaptic maturation, proteins selected for orthogonal validation, including Camk2a, Vamp2, Syp and Cplx1/2, exhibited steep increases in abundance after DIV11 (Fig. 1I). This pattern was representative of the broader synaptic proteome, with additional pre- and postsynaptic proteins, including Nrxn2, Syt1 and Gphn, and associated factors, Kif2a, Robo2, Ptprs, Septin3, displaying similar temporal trajectories (Fig. 1I). Immunostaining independently confirmed the increased abundance of Camk2a, Vamp2, Syp and Cplx1/2 at DIV35 relative to DIV11 (Fig. 1J). In line with the increase in synaptic protein expression, a proteomic time-course analysis demonstrated a progressive increase in the abundance of proteins involved in myelination and oligodendrocyte maturation^43^ (Fig. S1G), which was also confirmed by immunofluorescence (Fig. S1H). Together, these findings indicate that the organoids recapitulate the key molecular features of late-stage neural maturation, including coordinated synaptic maturation and myelination.

Having established the progressive maturation of the synaptic proteome during organoid development, we next examined how closely mature organoids recapitulate the *in vivo* synaptic proteome. In order to address this, we selectively isolated mature synaptic compartments for proteomic analysis. To enable synapse-specific enrichment, organoids were transduced with AAV-SypGreen (Synaptophysin coupled to a Green-fluorescent protein) at DIV16 and synaptosomes were prepared from DIV35 cultures, followed by fluorescence-activated synaptosome sorting (FASS) to isolate SypGreen-positive synaptosomes (Fig. 2A and 2B). Immunostaining of organoid sections demonstrated the colocalisation of SypGreen with either the inhibitory presynaptic marker Vgat and the excitatory presynaptic marker Vglut1 (Fig. 2C). Pixel-wise fluorescence intensity correlation, quantified using Pearson’s correlation coefficient, was significantly higher for Vglut1 than Vgat in both organoids and adult cortex, indicating a stronger correspondence between SypGreen and excitatory presynaptic labelling (Fig. 2D). Notably, transduction-mediated labelling of excitatory (Vglut1⁺) and inhibitory (Vgat⁺) synaptic termini in organoids yielded labelling efficiencies equivalent to those obtained in the cortex using a Syn1-cre driver line (Fig. 2D). This similarity in labelling efficiency may, at least in part, reflect the comparable fractional representation of excitatory (Vglut1⁺) and inhibitory (Vgat⁺) neuronal populations and their synaptic termini in organoids and the cortex. We next verified that the established Percoll synaptosome procedure^25,44^ reliably leads to a synaptosome-enriched fraction when using neuronal organoids as input material. Western blot analysis of the synaptosome fraction (F2/3) demonstrated enrichment of established synaptic proteins, including Stx1b, Syt12, Syp, Psd95, Homer1, Vamp2 and Bsn, relative to the starting homogenate, together with marked depletion of nuclear (H3), myelin (Mbp) and astrocytic (Gfap) markers, indicating enrichment of synaptic material while effectively removing major non-synaptic contaminants (Fig. 2E). Electron microscopy further confirmed the presence of structurally preserved synaptic structures within the F2/3 fraction, including both synaptosomes and synaptoneurosomes with identifiable pre- and postsynaptic compartments, synaptic vesicles and a postsynaptic density (PSD) (Fig. 2F).

**Figure 2.**
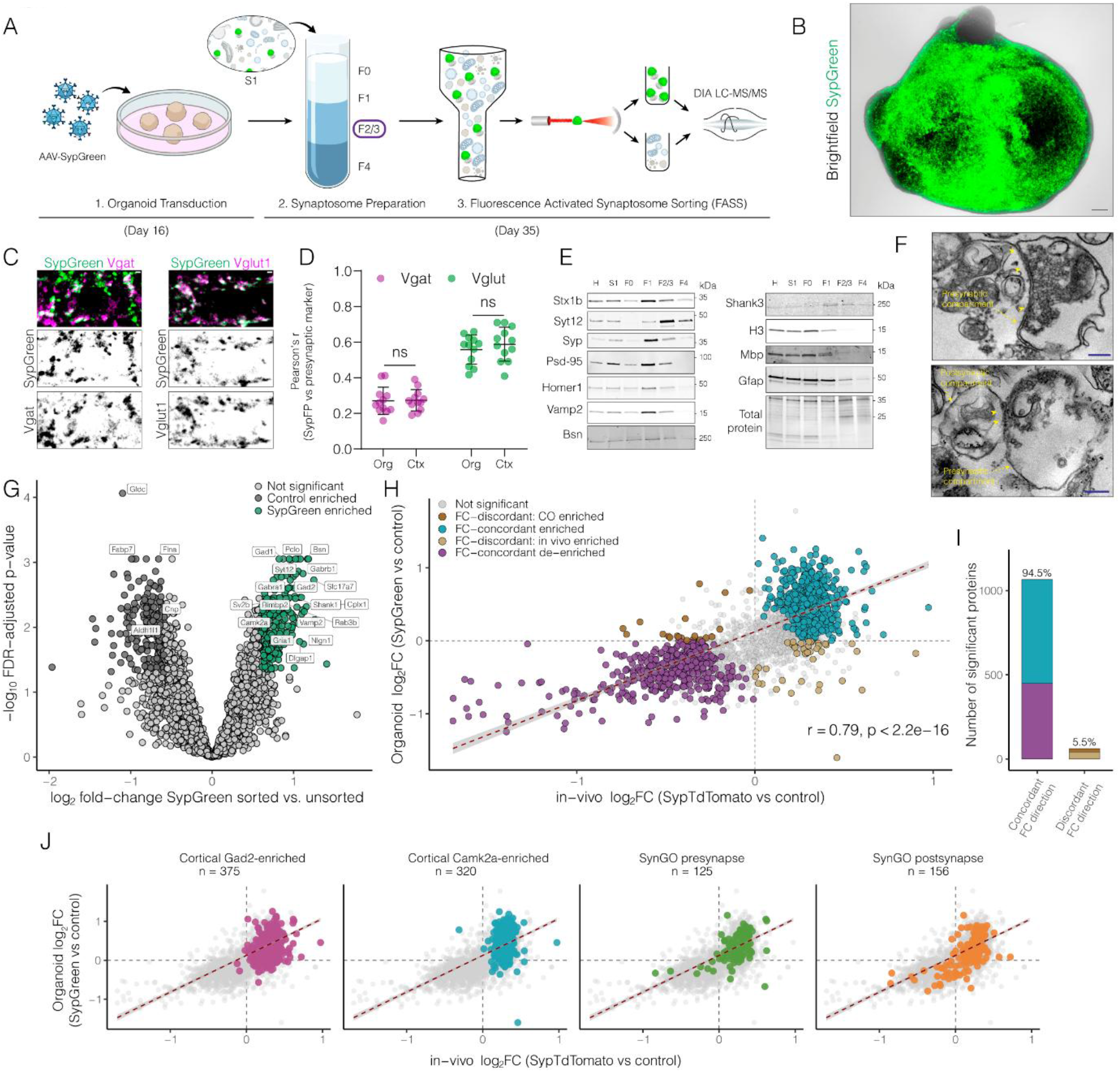
Mouse neural organoids recapitulate *in vivo* synaptic proteome enrichment patterns. (A) Overview of the synaptosome enrichment workflow. To selectively label presynaptic terminals, brain organoids were transduced with AAV-Synapsin–SypGreen (AAV-SypGreen) at DIV16 . At DIV35, organoids were homogenized, synaptosomes were isolated by discontinuous percoll density gradient centrifugation, and the synaptosome-enriched F2/3 fraction was subjected to fluorescence-activated synaptosome sorting (FASS) to isolate SypGreen-positive populations for quantitative DIA LC–MS/MS proteomic analysis. (B) Representative brightfield and fluorescence image of a DIV35 brain organoid following AAV-SypGreen transduction, demonstrating widespread neuronal expression of the synaptic reporter throughout the organoid. (C) Representative confocal images of SypGreen-expressing DIV35 organoids co-immunostained with the inhibitory presynaptic marker VGAT or the excitatory presynaptic marker VGLUT1. Images demonstrate robust overlap between SypGreen fluorescence and canonical presynaptic markers. Scale bars, 1 μm. (D) Quantification of colocalization between SypGreen and VGAT or VGLUT1 in organoid sections (Org) and cortex (Ctx) of Syn1-cre::CondSypTOM mice. Pearson’s correlation coefficients indicate comparable labeling specificity between organoid and cortical preparations, with no significant differences (ns). (E) Immunoblot analysis of the fractions obtained during synaptosome purification. Presynaptic proteins (Stx1b, Syt12, Syp, Vamp2, and Bsn) together with postsynaptic proteins (Psd95, Shank3 and Homer1) are enriched in the synaptosome-containing F2/3 fraction relative to homogenate (H), S1, F0, F1, and F4 fractions, confirming successful isolation of synaptic material. H3, Mbp and Gfap demonstrate depletion of nuclear, myelin, and astroglial contaminants, respectively. Total protein staining is shown as a loading control. (F) Representative transmission electron micrographs of synaptosomes and synaptoneurosomes from the F2/3 fraction. (Top) Representative presynaptic terminal containing numerous synaptic vesicles juxtaposed to an electron-dense postsynaptic density, with the postsynaptic compartment open. (Bottom) Representative synaptoneurosome comprising a presynaptic compartment containing synaptic vesicles and an adjacent resealed postsynaptic compartment. Yellow arrowheads indicate the postsynaptic density (PSD), and yellow dashed arrows delineate the pre- and postsynaptic compartments. Blue scale bars, 200 nm. (G) Volcano plot showing differential protein abundance between SypGreen-positive sorted synaptosomes and unsorted synaptosome preparations. Proteins significantly enriched in the SypGreen-positive fraction are highlighted in green, whereas proteins enriched in the control fraction are shown in dark grey. Representative synaptic proteins associated with neurotransmitter release, synaptic vesicles, and postsynaptic organization are annotated. (H) Comparison of differential protein abundance between organoid-derived SypGreen-positive synaptosomes and an independently generated *in vivo* cortical synaptosome dataset (SypTdTomato-positive versus control). Scatter plot demonstrates high correlation between datasets (Pearson’s r = 0.79, P < 2.2 × 10⁻¹⁶). Proteins are colored according to concordant enrichment, concordant depletion, or discordant regulation between datasets. Dashed lines indicate zero log₂ fold change. (I) Proportion of significantly regulated proteins exhibiting concordant or discordant differential abundance between organoid-derived and *in vivo* synaptosome datasets. The majority of regulated proteins (94.5%) display concordant fold-change direction, whereas only a small fraction (5.5%) exhibit discordant regulation. (J) Concordance analysis for biologically relevant protein subsets. Scatter plots show cortical GAD2-enriched proteins, cortical CAMK2a-enriched proteins, SynGO presynaptic proteins, and SynGO postsynaptic proteins overlaid on the global dataset. Each subset demonstrates strong agreement between organoid-derived and *in vivo* synaptosome enrichment profiles, indicating that fluorescence-assisted synaptosome sorting faithfully captures both inhibitory and excitatory synaptic proteomes as well as canonical pre- and postsynaptic molecular architectures.

Next, we performed quantitative proteomic analysis of FASS-isolated SypGreen⁺ synaptosomes from mature, spontaneously active organoids. FASS efficiently enriched fluorescently labelled synaptosomes, increasing the proportion of SynpGreen-positive particles around 7-fold compared to the input F2/3 fraction (Fig. S2A). We purified SypGreen+ synaptosomes from six independent biological replicates and quantified more than 4,000 protein groups across all mass spectrometry runs with similar protein abundance distributions between unsorted and sorted samples (Fig. S2B,C). Consistent with the selective isolation of a distinct synaptic population, principal component analysis clearly separated sorted and unsorted populations (Fig. S2D). Differential proteomic analysis confirmed that this shift reflected a significant enrichment of more than 750 protein groups in the SypGreen⁺ fraction, including canonical pre- and postsynaptic markers such as Gad1/2, Slc17a7, Camk2a, Shank1, Cplx1, Bsn, Ssyt12 and Vamp2, relative to the control fraction (Fig. 2G).

To assess the extent to which organoid-derived synapses reproduce the molecular architecture of mature cortical synapses, we compared their synaptic proteomic profiles with those obtained from *in vivo* cortical synaptosomes. Synaptic enrichment was concordant between the two preparations (Pearson’s r = 0.79), with >94% of significantly regulated proteins exhibiting a consistent directionality of enrichment (Fig. 2H,I). This concordance was evident for both inhibitory and excitatory neuronal markers, as well as SynGO-defined pre- and postsynaptic protein sets (Fig. 2J), indicating that organoid-derived synapses faithfully recapitulate the molecular composition of mature cortical synapses. To define the conserved molecular features underlying this concordance, we examined proteins that were consistently enriched or depleted following FASS in both datasets. Concordantly enriched proteins showed strong SynGO enrichment for canonical synaptic compartments, including presynaptic and postsynaptic structures, synaptic vesicles, active zone components, synaptic membranes and the synaptic cytoskeleton (Fig. S2E), which was mirrored by Gene Ontology enrichment for synapse, presynapse, cell junction and neuronal projection cellular components, together with biological processes involved in synaptic vesicle cycling, vesicle-mediated transport, chemical synaptic transmission and trans-synaptic signalling (Fig. S2F,G). In contrast, proteins consistently depleted by FASS were dominated by mitochondrial cellular components and associated metabolic and translational processes (Fig. S2I,J) and exhibited a modest enrichment for SynGO synaptic annotations, limited to pre-/postsynaptic ribosomes (Fig. S2H). Together, these findings demonstrate that the synaptic proteome of mouse cortical organoids closely recapitulates the molecular composition of mature *in vivo* cortical synapses.

## Discussion

Cerebral organoids provide experimental access to human neuronal tissue and non-model organisms that are otherwise inaccessible, making them a powerful platform for studying synapse biology^6,26^. Because synaptic dysfunction is a central feature of many neurodevelopmental and neurodegenerative disorders, and synaptic proteins represent important therapeutic targets^16,17^, it is essential to establish whether organoid synapses can faithfully recapitulate their *in vivo* counterparts. Recent studies have begun to define the molecular composition of human organoid synapses, revealing substantial similarities with developing brain tissue^29^. However, because human neural development proceeds over extended timescales^30–33,35^, it has remained unclear whether the incomplete synaptic maturation observed in these systems reflects intrinsic developmental timing or limitations imposed by *in vitro* culture.

By leveraging the accelerated developmental trajectory of mouse organoids^36,45^ together with a matched adult mouse cortical reference dataset, our study demonstrates that mouse forebrain organoids possess a mature synaptic proteome that closely recapitulates the molecular architecture of adult cortical synapses. By combining developmental proteomics with fluorescence-activated synaptosome sorting and quantitative mass spectrometry, we show that synaptic maturation in organoids is accompanied by the coordinated expression of canonical pre- and postsynaptic proteins and that purified synaptosomes exhibit high molecular similarity to those from the adult cortex. These findings show that, despite the absence of a native brain environment, organoid-derived neurons can establish a complex synaptic molecular organization that resembles mature *in vivo* synapses. We thus established that, given sufficient developmental progression, organoid systems are capable of fully establishing the molecular features that characterize mature cortical synapses.

Because synapse-type-specific proteomics provides a direct means of quantifying the molecular machinery underlying the function of genetically defined synapse types^46,47^, it is a powerful approach for probing synaptic protein alterations underlying neurological diseases^48–50^. By demonstrating the feasibility of isolating and quantitatively profiling organoid-derived synaptosomes, this work establishes a framework for applying such analyses to human organoid models of neurodevelopmental and neurodegenerative disease^9,12,35,51,52^. The combination of organoid models with FASS-based synaptosome purification will facilitate mechanistic studies of synaptic pathology and enable the evaluation of therapeutic interventions in genetically tractable systems.

---

**Figure Sup 1.**
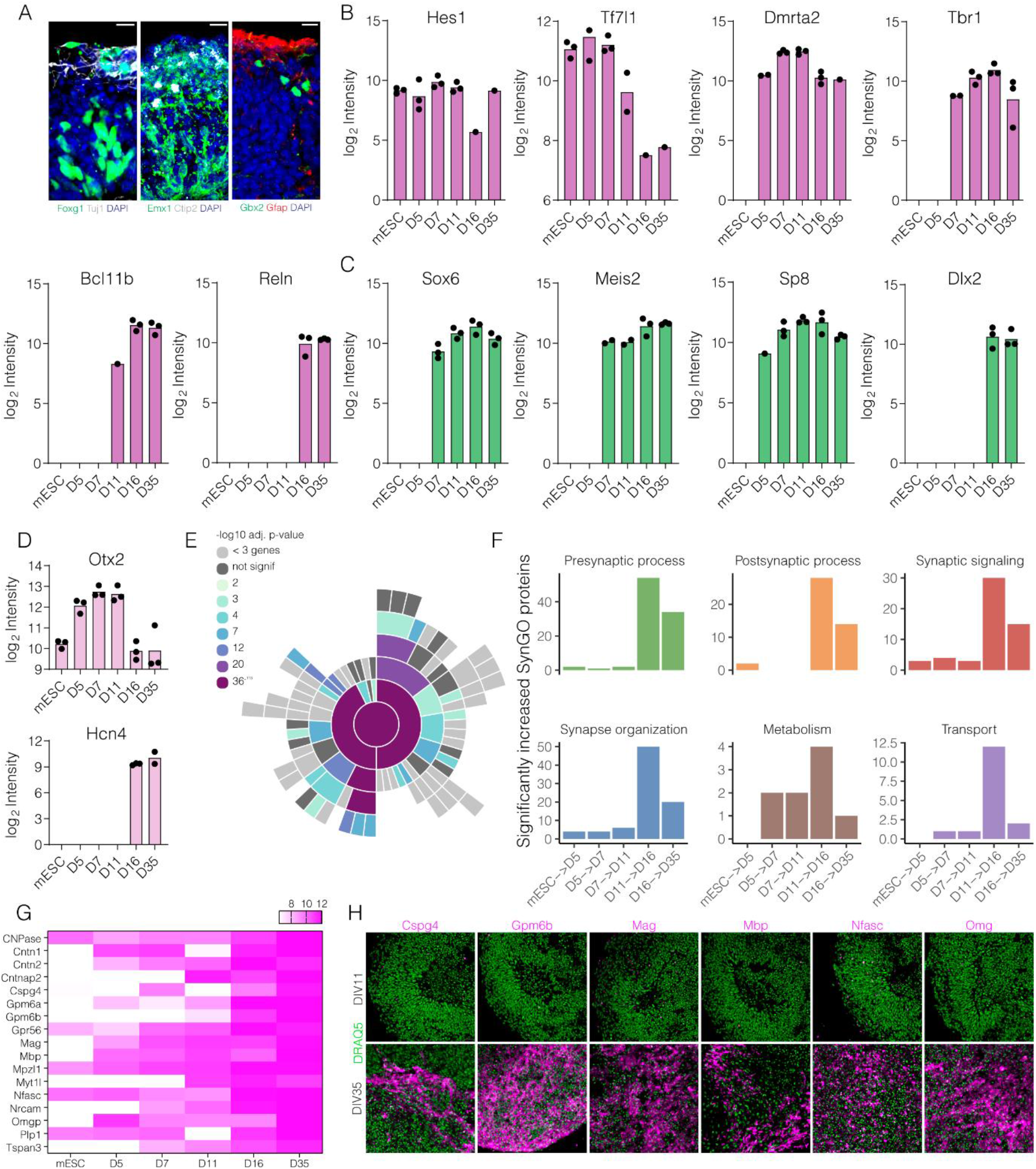
Identity marker expression and synaptic gene ontology dynamics during organoid maturation. (A) Representative immunofluorescence images demonstrating the acquisition of forebrain and cortical identity in mouse brain organoids. Organoids express the forebrain marker Foxg1 together with the neuronal marker Tuj1, the dorsal cortical progenitor marker Emx1 and deep-layer cortical neuron marker Ctip2, and the thalamic marker Gbx2 together with the astrocytic marker Gfap. Nuclei were counterstained with DAPI. Scale bars, 20 μm. (B) Quantitative proteomic analysis of dorsal forebrain developmental markers across organoid differentiation. Log₂ protein abundances are shown for the neural progenitor marker Hes1, the corticothalamic neuron marker Tbr1, and additional dorsal telencephalic transcription factors Tf7l11 (Tcf7l1), Drmta2, Bcl11b (Ctip2), and Reln. Data demonstrate the progressive emergence of cortical neuronal identity during maturation. (C) Temporal protein abundance profiles of ventral telencephalic and interneuron-associated transcription factors. Expression of Sox6, Meis2, Sp8, and Dlx2 indicates the presence of ventral forebrain-derived neuronal populations within organoids and their maturation over time. (D) Quantitative proteomic analysis of thalamic markers. Otx2 expression is highest during early differentiation and decreases with maturation, whereas Hcn4 expression emerges at later developmental stages. (E) SynGO cellular component enrichment analysis of proteins exhibiting significant temporal increases during organoid maturation. Circular enrichment plot demonstrates progressive enrichment of proteins associated with presynaptic and postsynaptic compartments, synaptic vesicles, active zone, postsynaptic density, dendritic spines, and additional synaptic structures. Color intensity indicates enrichment significance (−log₁₀ adjusted P value). (F) Distribution of significantly increased SynGO-annotated proteins across major functional categories during consecutive developmental transitions. The largest increases occur during the DIV11-to-DIV16 and DIV16-to-DIV35 transitions and are associated with presynaptic processes, postsynaptic processes, synaptic signaling, synapse organization, metabolism, and transport, indicating extensive synaptic maturation during late-stage organoid development. (G) Heatmap showing temporal expression profiles of representative oligodendrocyte- and myelin-associated proteins identified by quantitative proteomics, including Cnp, Cntn1, Cntn2, Cntnap2, Cspg4, Gpm6a, Gpm6b, Gpr56, Mag, Mbp, Mpzl1, Myt1, Nfasc, Nrcam, Omg, Plp1, and Tspan3. Color scale represents log₂ protein abundance. (H) Immunofluorescence validation of oligodendrocyte lineage and myelin-associated protein expression in D11 and D35 brain organoids. Representative images show expression of Cspg4, Gpm6b, Mag, Mbp, Nfasc, and Omg together with the nuclear counterstain DRAQ5 (green), demonstrating the emergence of oligodendrocyte-associated cell populations and myelin-related proteins during organoid maturation. Scale bars, 20 μm. Data represent independent biological replicates analyzed by quantitative DIA LC–MS/MS. Protein abundances are displayed as log₂-transformed intensities.

**Figure Sup 2.**
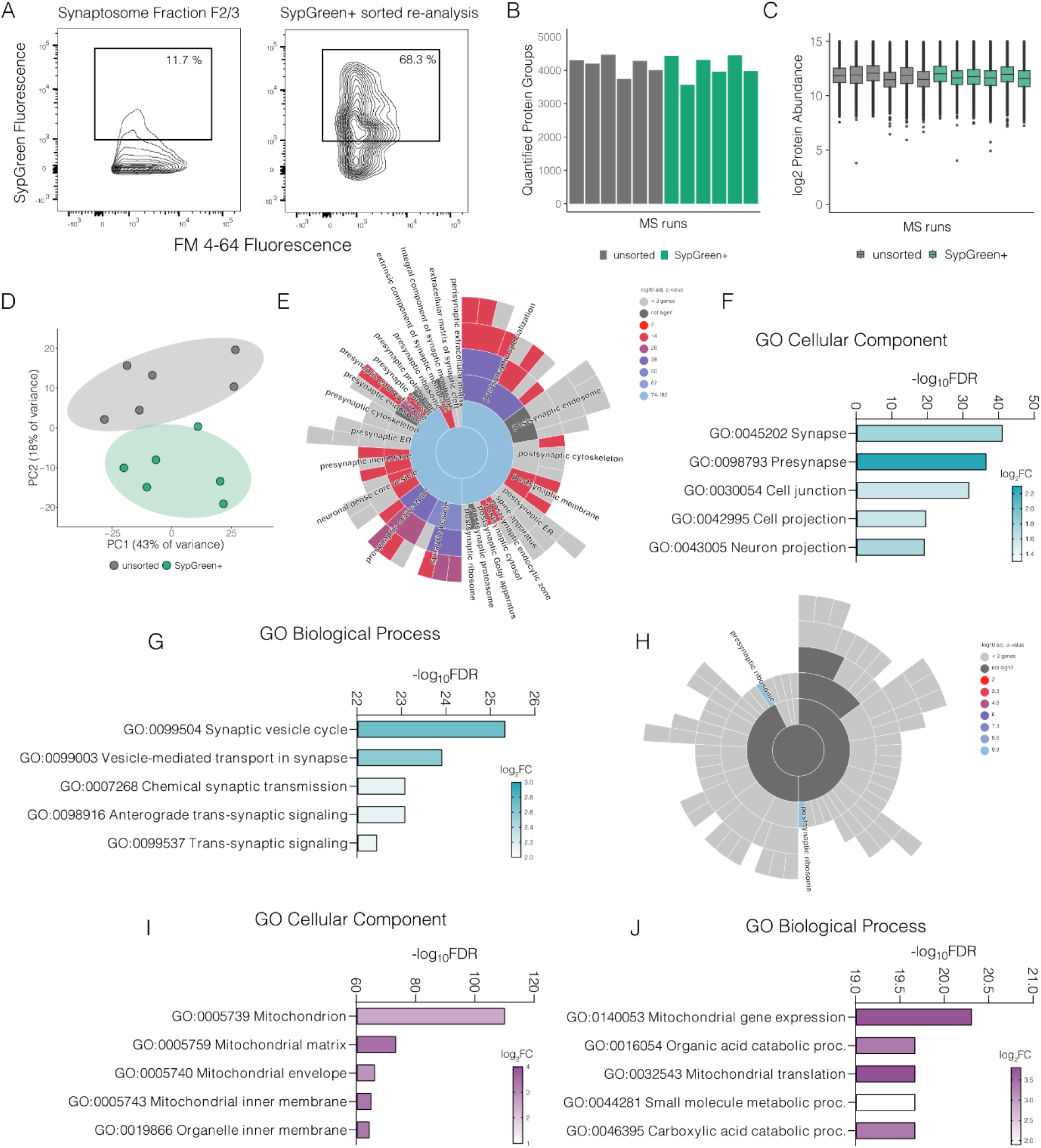
Synaptic proteomes enriched by FASS exhibit conserved molecular signatures across organoids and cortex. (A) Representative flow cytometry plots illustrating fluorescence-assisted synaptosome sorting (FASS). Left, gating strategy applied to the synaptosome-enriched F2/3 fraction based on FM4-64 membrane labeling and SypGreen fluorescence. Right, re-analysis of the sorted population demonstrating enrichment of SypGreen-positive synaptosomes following sorting. (B) Number of quantified protein groups identified in unsorted synaptosome preparations and SypGreen-positive sorted synaptosome samples across individual DIA LC–MS/MS runs, demonstrating consistent proteome depth between experimental groups. (C) Distribution of protein abundances across all mass spectrometry runs shown as boxplots of log₂-transformed protein intensities. Comparable abundance distributions indicate high analytical reproducibility and effective normalization between unsorted and SypGreen-positive samples. (D) Principal component analysis (PCA) of quantitative proteomic datasets. Unsorted and SypGreen-positive synaptosome samples separate along the principal components, indicating distinct proteomic compositions following fluorescence-assisted sorting while maintaining tight clustering of biological replicates. (E) SynGO cellular component enrichment analysis of proteins significantly enriched in SypGreen-positive synaptosomes. Circular enrichment plot demonstrates strong overrepresentation of canonical presynaptic structures, including synaptic vesicles, presynaptic active zone, presynaptic membrane, presynaptic cytoskeleton, synaptic vesicle membrane, synaptic vesicle lumen, and presynaptic endosome, together with enrichment of additional synaptic compartments. Color intensity reflects enrichment significance (−log₁₀ adjusted P value). (F) Gene Ontology (GO) Cellular Component enrichment analysis of proteins enriched in SypGreen-positive synaptosomes. The most significantly enriched categories include synapse, presynapse, cell junction, cell projection, and neuron projection. Bar color indicates mean log₂ fold-change of proteins contributing to each enriched term. (G) GO Biological Process enrichment analysis of proteins enriched in SypGreen-positive synaptosomes. Significantly enriched processes include synaptic vesicle cycle, vesicle-mediated transport in synapse, chemical synaptic transmission, anterograde trans-synaptic signaling, and trans-synaptic signaling, consistent with selective enrichment of functional presynaptic machinery. (H) SynGO cellular component enrichment analysis of proteins significantly depleted following SypGreen sorting. Depleted proteins are predominantly associated with non-synaptic compartments, postsynaptic ribosomes and other intracellular structures, indicating efficient removal of contaminating cellular material from the sorted synaptosome preparation. (I) GO Cellular Component enrichment analysis of proteins depleted in the SypGreen-positive fraction. Mitochondrial components, including mitochondrion, mitochondrial matrix, mitochondrial envelope, mitochondrial inner membrane, and organelle inner membrane, are significantly overrepresented among depleted proteins, demonstrating reduced contamination by mitochondrial proteins after FASS. (J) GO Biological Process enrichment analysis of proteins depleted in SypGreen-positive synaptosomes. Enriched terms include mitochondrial gene expression, mitochondrial translation, organic acid catabolic process, small molecule metabolic process, and carboxylic acid catabolic process, consistent with selective depletion of metabolic and mitochondrial pathways relative to the enriched synaptic proteome. Data represent independent biological replicates processed by DIA LC–MS/MS. Differential abundance and enrichment analyses were performed as described in the Methods. Colour scales represent mean enrichment significance (−log₁₀ adjusted P value,) or log₂ fold change, as indicated in each panel. The unsorted synaptosome proteome was used as background for GO enrichment analyses.

## Methods

### mESC culture

ES-E14Tg2A (ATCC, CRL-1821) and R1/E (ATCC, SCRC-1036) cells were maintained as feeder-free cultures on gelatin-coated plates in Serum/LIF medium (DMEM+Glutamax, 0.001% 2-mercaptoethanol, MEM amino acids and 10% FCS, 2000 U/mL ESGRO LIF), as previously described^53^. Cells were grown at 37°C in 5% CO_2_, passaged weekly with TrypLE Express and regularly mycoplasma tested.

### Neural organoid generation

Neural organoids were generated following an adapted unguided cerebral organoid protocol^37,41^ and using the STEMdiff Cerebral Organoid kit (STEMCELL Technologies, 08570) reagents. Briefly, mESCs were dissociated and seeded into low-attachment culture plates in embryoid body (EB) medium to promote embryoid body formation (Day 0). EBs were cultured for 5 days until transfer to neural induction (NI) medium to promote the induction of neuroepithelial identity. On day 7, organoids were embedded in Matrigel Growth Factor Reduced (MG GFR) to induce neuroepithelial polarity reversal and were transferred to expansion medium. From day 11, organoids were maintained in maturation medium under continuous orbital agitation to enhance nutrient and oxygen diffusion and promote tissue growth. Beginning on day 16, the culture medium was supplemented with brain-derived neurotrophic factor (BDNF, 20 ng/ml) and neurotrophin-3 (NT-3, 20 ng/ml) to support neuronal differentiation and maturation. Organoids were cultured until Day 35, at which point they displayed hallmarks of neuronal maturity and were used for downstream functional analyses of synapses.

### Immunofluorescence

Organoids were fixed in 4% PFA in PBS-MC+Sucrose (PBS pH 7.4, 1 mM MgCl2, 0.1 mM CaCl2, 4% sucrose) overnight at 4℃. Fixation was stopped by 3 washes in PBS-MC, tissues were allowed to sink in 30% sucrose overnight for cryoprotection, embedded in Tissue-Tek O.C.T. compound (Sakura) and cryosectioned at 20-25 μm. Sections were blocked and permeabilized with 0.5 % Triton X-100, 4% goat serum in PBS (pH 7.4) for 2h and then incubated overnight with primary antibodies in 4% goat serum in PBS (pH 7.4) at the following dilutions: rb gp anti-Bsn (SYSY, 141 318, 1:200), rb anti-Camk2a (Abcam, ab52476, 1:500), rb anti-Cspg4 (Proteintech, 31623-1-AP, 1:200), rt anti-Ctip2 (Abcam, ab18465, 1:500), rb anti-Cplx1/2 (SYSY, 122 102, 1:200), rb anti-Dlx2 (Proteintech, 26244-1-AP, 1:200), rb anti-Emx1 (Proteintech, 55032-1-AP, 1:200), rb anti-Foxg1 (Proteintech, 85930-1-RR, 1:200), ms anti-Gad65/67 (Enzo, ADI-MSA-225-E, 1:200), rb anti-Gbx2 (Proteintech, 21639-1-AP, 1:200), ms anti-Gfap (Abcam, ab10062, 1:500), gp anti-Gphn (SYSY, 147 318, 1:200), rb anti-Gpm6b (Proteintech, 26122-1-AP, 1:200), ms anti-HuC/D (Thermo Fisher, A21271, 1:1000), rb anti-Mag (Proteintech, 14386-1-AP, 1:200), chk anti-Map2 (Abcam, ab5392, 1:1000), rb anti-Mbp (Abcam, ab40390, 1:200), rb anti-Nfasc (Proteintech, 26351-1-AP, 1:200), ms anti-NeuN (Millipore,MAB377, 1:200), rb anti-Omg (Proteintech, 12701-1-AP, 1:200), rb anti-Pax6 (Abcam, ab195045, 1:200), ms anti-pH3 (Abcam, ab14955, 1:500), anti-Psd95 (SYSY, 124 003, 1:200), ms anti-Syp (Sigma-Aldrich, S5768, 1:500), rb anti-Tbr2/Eomes (Proteintech, 83945-5-RR, 1:200), ms anti-Tuj1 (Sigma-Aldrich, T8578, 1:1000), rb anti-Vamp2 (SYSY, 104 202, 1:200), gp anti-Vgat (SYSY, 131 004, 1:200), rb anti-Vglut1 (Abcam, ab77822, 1:200). Samples were incubated with DAPI (1 µg/mL) and 405/488/568/647 Alexa Fluor-conjugated goat secondary antibodies (Thermo Fisher, DF=1:500) for 30 min at room temperature. The slides were coverslipped with Aqua-Poly/Mount.

### Microscopy

Images were acquired on either a Zeiss LSM780 or 880 inverted confocal microscopes using LD LCI Plan-Apochromat 63x/1.2 Imm Corr DIC M27, Plan-Apochromat 40x/1.3 Oil DIC UV-IR M27 and Plan-Apochromat 20x/0.8 M27 objectives.

### Viral transduction

Viral particles for the pssAAV-2-hSyn1-GCaMP6s_2A_NLS_dTomato-WPRE-hGHp(A) (p157) and pssAAV-2-hSyn1-mSyp1_CaMPARI2(F391W, L398V, no tags)-WPRE-hGHp(A) (p515; hereby referred to as SypGreen in its non-photoconverted form) constructs were obtained from the viral vector facility of the University of Zurich in the AAV-DJ serotype. On day 16 of the protocol, 3-6 organoids were transduced with a number of viral particles ranging between ∼15-30×10^6^ vg.

### Calcium imaging

On day 16 of the protocol, organoids were transduced with pssAAV-2-hSyn1-GCaMP6s_2A_NLS_dTomato-WPRE-hGHp(A) (p157). Calcium imaging was performed at 30-40 days *in vitro* using a Zeiss LSM 880 confocal microscope with a 20×/0.8 air objective, acquiring at 2-3 fps. Imaging was conducted in culture maturation medium on a 37 °C, 5% CO₂ incubated stage.

### Synaptosome preparation

Synaptosomes were isolated following the protocol described by Westmark et al.^44^ with minor modifications^25^. Organoids were homogenized in gradient medium (GM; 0.25 M sucrose, 5 mM Tris-HCl, 0.1 mM EDTA supplemented with Calbiochem Protease Inhibitor Cocktail III) using a glass Dounce homogenizer. The homogenate was centrifuged at 1,000 × g for 10 min at 4°C to obtain the supernatant (S1). The S1 fraction was layered onto a discontinuous Percoll density gradient consisting of 23%, 10%, and 3% Percoll prepared in GM buffer. Gradients were centrifuged at 32,500 × g for 5 min at 4°C using maximum acceleration and minimum deceleration in a Beckman Coulter JA-25.50 rotor mounted in an Avanti J-26S XPI centrifuge (Beckman Coulter). Following centrifugation, four fractions (F0, F1, F2/3, and F4, from top to bottom) were collected.

### Fluorescence-activated synaptosome sorting

Fluorescence-activated synaptosome sorting (FASS) was performed as previously described^25,39^. F2/3 fractions from the synaptosome preparation were diluted in the GM buffer and 1.5mg/ml membrane dye (FM4-64, Thermo Fisher) was added. Synaptosomes were analyzed and sorted on a FACSAria Fusion (BD Biosciences) running FACSDiva, equipped with a 70mm Nozzle and the following settings: 488nm laser (for FM4-64 and SypGreen), sort precision (0-16-0), FSC (317 V), SSC (488/10 nm, 370V), GFP ‘‘SypGreen’’ (530/30 nm, 535V), PerCP ‘‘FM4-64’’ (695/40 nm, 470), thresholds (FSC = 200, FM4-64 = 700). Samples were analyzed and sorted at approx. 20,000 events/s and a flow rate of < 3. Doublet particles were excluded based on SSC-H and SSC-W. For each sorted sample (SypGreen+) as well as the matching control sample (FM+) we sorted 5 Mio particles.

### Synaptosome processing for MS

Sorted synaptosomes and control particles were filtered onto Whatman glass microfiber filters (GF/F, Cytiva) and stored at -80°C until further processing as described before^25^. For filtration, a miniaturized custom-built apparatus was constructed. Synaptosomes were sorted directly into a 15ml tube (Protein LoBind, Eppendorf) that was connected through tubing (from an infusion set) to a luer connector unit with a 4mm diameter filter fixed between the male and female luer connector units. The tubing and filtration unit was cooled on ice and negative pressure was applied for filtration. Synaptosome samples were solubilized in 20ul TEAB buffer (50mM Triethylammonium bicarbonate, 1mM CaCl, 0.05μg trypsin LysC and 0.05μg trypsin) and digested overnight at 37°C. 30μl acetonitrile was added and the samples were centrifuged 10 min at 16,000 g. The supernatant was filtered through ZipTip pipette tips by centrifugation for 1 min at 2,000g and then 50μl of 50% acetonitrile in MS-grade water was added to the filter and both steps were repeated. Samples were dried in a vacuum centrifuge and stored at - 20°C until Liquid chromatography–tandem mass spectrometry (LC-MS/MS) analysis. Organoid time series samples were prepared using an SP3 protocol^54,55^ as previously described^56^. In brief, organoids were washed in DPBS+Ca^2+^/Mg^2+^ (Invitrogen, 14040141) and lysed in RIPA buffer (Invitrogen, 89900). 0.5 µl magnetic, carboxylate-modified beads (1:1 mix of Sera-Mag SpeedBeads (GE Healthcare, 45152105050250 & 65152105050250; pre-washed in MS-grade water three times) were added to the samples. Protein binding was facilitated by addition of 100% MS-grade ethanol to a final concentration of 50% ethanol and shaking for 5 min at RT at 1000 rpm. Additional incubation was carried out for 15 min at RT without agitation. Beads were then washed three times with 100 µl 80% ethanol using a magnetic rack. Proteins were digested in 20 µl digest buffer (50 mM triethylammonium bicarbonate) with 0.1 µg MS-grade trypsin (Promega) in combination with 0.1 µg LysC (Wako) at 37°C shaking at 1000 rpm overnight. Peptides were desalted using C18 zip tips (cat. no. ZTC18S960, Millipore), dried by vacuum centrifugation and stored at −20°C until LC-MS analysis.

### LC-MS/MS analysis

Dried peptides were reconstituted in 5% acetonitrile and 0.1% formic acid with iRT peptide standard (Biognosys, 1:100) additionally spiked into synaptosome samples. Peptides were loaded onto a PepMap 100 C18 trapping column (20 mm × 75 μm, 3 μm particle size; Thermo Fisher Scientific) and separated on a C18 analytical column with integrated emitter (50 cm × 75 μm, 1.7 μm particle size; CoAnn Technologies) using an UltiMate 3000 RSLCnano system (Thermo Fisher Scientific) coupled to a nanoFlex source (2000 V, Thermo Fisher Scientific). The analytical column was maintained at 55 °C throughout. Trapping was performed for 6 min at 6 μL/min using 2% acetonitrile and 0.05% trifluoroacetic acid. Peptides were separated at 250 nL/min using a 120 min non-linear gradient of buffer A (0.1% formic acid in water) and buffer B (80% acetonitrile, 0.1% formic acid)^57^. MS analysis was performed on a Fusion Lumos Orbitrap mass spectrometer (Thermo Fisher Scientific) operated in positive-ion DIA mode. Full MS scans were acquired at 120,000 resolution over an m/z range of 350–1650, with an AGC target of 125% and a maximum injection time of 100 ms. DIA scans were acquired using 40 isolation windows with HCD fragmentation (collision energy 27%), a resolution of 30k, an AGC target of 2000%, and dynamic maximum injection time^57^.

### Processing of LC-MS/MS Data

LC-MS/MS samples were analyzed with Spectronaut version 19 (Biognosys) followed by statistical analysis using the MSstats R package^58–60^. Targeted data extraction of DIA-MS acquisitions was performed in DirectDIA mode with default settings using the UniProtKB/Swiss-Prot database for *Mus musculus* (retrieved in 2023), a database consisting of common mass spectrometry contaminants and the transgenic constructs. The proteotypicity filter “only protein group specific” was applied. Extracted features were exported from Spectronaut for statistical analysis with MSstats 4 using default settings. Briefly, for each protein, features were log-transformed and fitted to a mixed effect linear regression model for each sample. For significance testing, contaminants were filtered out, all values below 5 were set to NA and we required a minimum of 4 features (combination of precursor and fragment ion) and 8 measurements per protein per condition. The model estimated fold change and statistical significance for all conditions. The Benjamini–Hochberg method was used to account for multiple testing and p-value adjustment was performed on all proteins that met the fold-change cutoff. The mass spectrometry proteomics data have been deposited to the ProteomeXchange Consortium via the PRIDE partner repository with the dataset identifier PXD084797.

### SDS-PAGE and immunoblotting

For immunoblotting, synaptosomes were lysed by adding lysis buffer (8 M urea, 10% SDS, 10% sodium deoxycholate, and 5% Triton X-100 in water) at a 1:5 (v/v) ratio of lysis buffer to synaptosome fraction, followed by incubation at 75°C for 5 min. Protein concentrations in each fraction were measured using the Precision Red advanced protein assay (Cytoskeleton, Inc). For each sample, a volume corresponding to a set protein amount was supplemented with 10X SDS sample buffer (500 mM Tris pH 6.8, 25% SDS and 2% bromophenol blue in 70% glycerol-30% dH2O), NuPAGE Sample Reducing Agent (10X) (Thermo Fisher, NP0004) and distilled water to an equal final volume. Samples were denatured and reduced at 85°C for 5 minutes and run on Novex 4-20% Tris-Glycine, Novex 12% Bis-Tris mini gels and 4-20% BioRad midi gels. After electrophoresis, proteins were semi-dry transferred onto 0.2 μm nitrocellulose membranes (BioRad, 1704159). Equal loading and even transfer were then assessed by Revert 700 total protein stain (LI-COR, 926-11011). The membranes were destained, blocked for 1h at room temperature in Intercept (TBS or PBS) blocking buffer (LI-COR, 927-60001, 927-70001) and probed with primary antibodies overnight at 4°C. The next day the membranes were developed with fluorescently labeled secondary antibodies on a LI-COR OdysseyF system. The following primary antibodies were used at a 1:1000 DF: ms anti-Bsn (Enzo, ADI-VAM-PS003-F), rb anti-Gfap (Abcam, ab7260), rb anti-H3 (Abcam, ab1791), rb anti-Homer1 (SYSY, 160 002), ms anti-Mbp (Abcam, ab62631), ms anti-Psd95 (Biozol, GTX22723), rb anti-Shank3 (Cell Signalling, 64555), ms anti-Syp (Sigma-Aldrich, S5768), rb anti-Stx1b (SYSY, 110 403), rb anti-Syt12 (SYSY, 299 003), rb anti-Vamp2 (SYSY, 104 202).

### Electron microscopy

Electron microscopy (EM) analysis of unsorted synaptosomes was conducted as described in Sebring et al.^61^, with chemicals from Sigma Aldrich and Plano GmbH (Pioloform), unless otherwise specified. A volume of ∼2 ml of F2/3 fractions were collected in 5 ml Eppendorf tubes and fixed with EM grade glutaraldehyde (final concentration 2.5%) for 30 minutes on ice and, subsequently, diluted in ∼3 ml PBS to stop fixation. The fixed synaptosomes were pelleted using 5427R Eppendorf centrifuge at ∼4000g for 1 hour at 4°C. Then, the supernatant was aspirated carefully and the pellet was resuspended in 5% low-melt agarose in MilliQ water. Samples were washed 3 times in 0.1M cacodylate buffer (Serva) and incubated with 1% osmium tetroxide (Science Services) for 1 hour. After 3x 5 minute wash in 0.1M cacodylate buffer, then 3x 3 minute wash in MilliQ water, samples were incubated for 10 minutes in 1% uranyl acetate (Serva). Samples were then washed 3x 5 minutes with MilliQ water, then dehydrated using an ascending ethanol series (30%, 50%, 70%, 90%, 2x 100%) for 5 minutes each, followed by 2x 5 minute incubation in propylene oxide (Merck). Infiltration was performed with a mixture of 50% EPON (Sigma Aldrich), and 50% propylene oxide for 30 minutes at room temperature. Samples were then incubated in 100% EPON overnight at room temperature. For sample embedding, excess agarose was trimmed and samples placed in a silicon mold (Plano) with fresh EPON and polymerized for at least 48 hours at 60°C. Ultrathin sections of 70nm were cut using a Leica Reichert Ultracut S Ultramicrotome with a DiATOME ultra 45° diamond knife and transferred to a single slot grid (TEM, grid, 2x1 mm, slit, Cu, Science Services, #G2010-Cu) previously coated with 1.1% Pioloform in chloroform. Grids were stored in grid boxes (Science Services, G71135-01) until imaging. TEM imaging was performed with a Zeiss Leo 912 AB Omega and a Sharp Eye TRS (2x2k) CCD camera. ImageSP was used to control the CCD camera and to make tilescans with 20% overlap and imaging carried out at 120 kEV.

### Statistical analysis

For microscopy data comparisons between two groups were made using unpaired two-tailed t-tests. Statistical significance was set at p≤0.05, with a minimum of three biological replicates. For statistical analysis and figure design GraphPad Prism, R, Adobe Illustrator, Biorender.com and FigureLabs were used.

## Acknowledgments

We thank the MPIBR Imaging and Proteomics Facilities for assistance with data acquisition, in particular Sarah Candlish for EM data collection. We are grateful to members of the Schuman lab for helpful discussions, comments, and feedback, and to Nicole Fürst, Christina Thum, Bastian Krause, Susanne tom Dieck, Anita Kulak and Thomas M. Blok for technical support. This work was supported by a long-term fellowship from the Federation of European Biochemical Societies (FEBS) and a HESSEN HORIZON Marie Skłodowska-Curie fellowship to Stefano L. Giandomenico; grants from the Swiss National Science Foundation (SNSF; P2EZP3_191820, P400PB_199288 and PZ00P3_216106), the Hirschmann Stifung and Synapsis Foundation (2024-CDA01) to Marc van Oostrum; and funding from the Max Planck Society and the European Union (ERC, DiverseSynapse, grant 101054512) to Erin M. Schuman.

## Author contributions

S.L.G., M.v.O., and E.M.S. conceived the study. S.L.G. established and optimized the differentiation protocol; generated and maintained organoids; performed biochemical and immunofluorescence experiments; prepared samples for EM; and analyzed the data. M.v.O. generated and maintained organoids and performed FASS, proteomics sample preparation, data acquisition and analysis. Q.W. prepared and ran samples for proteomics. I.D.C. established and performed EM sample preparation. J.D.L. oversaw the proteomics data acquisition. S.L.G., M.v.O., and E.M.S. wrote the manuscript.

